# Neural speech tracking benefit of lip movements predicts behavioral deterioration when the speaker’s mouth is occluded

**DOI:** 10.1101/2023.04.17.536524

**Authors:** Patrick Reisinger, Marlies Gillis, Nina Suess, Jonas Vanthornhout, Chandra Leon Haider, Thomas Hartmann, Anne Hauswald, Konrad Schwarz, Tom Francart, Nathan Weisz

**Author notes:** Shared last authorship (Tom Francart, Nathan Weisz).

## Abstract

Observing lip movements of a speaker is known to facilitate speech understanding, especially in challenging listening situations. Converging evidence from neuroscientific studies shows enhanced processing of audiovisual stimuli. However, the interindividual variability of this visual benefit and its consequences on behavior are unknown. Here, we analyzed source-localized magnetoencephalographic (MEG) responses from normal-hearing participants listening to audiovisual speech with or without an additional distractor speaker. Using temporal response functions (TRFs), we show that neural responses to lip movements are, in general, enhanced when speech is challenging. After conducting a crucial control for speech acoustics, we show that lip movements effectively contribute to higher neural speech tracking, particularly when a distractor speaker is present. However, the extent of this visual benefit varied greatly among participants. Probing the behavioral relevance, we show that individuals who benefit more from lip movement information in terms of neural speech tracking, show a stronger drop in performance and an increase in perceived difficulty when the mouth is occluded by a surgical face mask. By contrast, no effect was found when the mouth was not occluded. We provide novel insights on how the benefit of lip movements in terms of neural speech tracking varies among individuals. Furthermore, we reveal its behavioral relevance by demonstrating negative consequences for behavior when visual speech is absent. Our results also offer potential implications for future objective assessments of audiovisual speech perception.

## Introduction

Face masks are an important tool in preventing the spread of contagious diseases such as COVID-19 (e.g. Chu et al., 2020; Suñer et al., 2022). However, as many have subjectively experienced first hand, the use of face masks also impairs speech perception, and not only by attenuating sound. More importantly, they occlude facial expressions, such as lip movements (e.g. Brown et al., 2021; Rahne et al., 2021), that provide visual information for a relevant speech stream. This is particularly critical when speech is challenging, such as in the classic cocktail party situation, where multiple conversations are happening simultaneously (Cherry, 1953). In such situations, the brain separates auditory information of interest from competing input (McDermott, 2009). Ideally, visual information is available to support this process, with numerous studies demonstrating that visual speech features enhance the understanding of degraded auditory input (e.g. Grant & Seitz, 2000; Remez, 2012; Ross et al., 2007; Sumby & Pollack, 1954). This concept is known as inverse effectiveness (Meredith & Stein, 1983; van de Rijt et al., 2019). Among visual speech features, lip movements are the most important, playing a crucial role in the perception of challenging speech (Erber, 1975; Peelle & Sommers, 2015). This is especially intriguing given the substantial interindividual differences in lip-reading performance among normal, as well as hearing-impaired, populations (Suess, Hauswald, Zehentner, et al., 2022; Summerfield et al., 1992). Despite our imperfect lip-reading abilities, the human brain effectively uses lip movements to facilitate the perception of challenging speech, with the neural mechanisms and regions involved still under debate (Ross et al., 2022; Zhang & Du, 2022).

Previous studies have shown beneficial effects of visual speech on the representation of speech in the brain. An MEG study by Park et al. (2016) showed enhanced entrainment between lip movements and speech-related brain areas when congruent audiovisual speech was presented. Other studies have shown that the incorporation of visual speech enhances the ability of the brain to track acoustic speech (Crosse et al., 2015; Crosse, Liberto, et al., 2016; Golumbic et al., 2013). Interestingly, when silent lip movements are presented, the brain also tracks the unheard acoustic speech envelope (e.g. Hauswald et al., 2018) or spectral fine details (Suess, Hauswald, Reisinger, et al., 2022). Despite these findings, two questions remain unanswered: First, it is unknown how individuals vary in their benefit of lip movements at the neural level. Given the aforementioned interindividual differences in lip-reading performance, a high degree of variability could also be expected here. Importantly, lip movements are correlated with acoustic speech features (Chandrasekaran et al., 2009), so it is essential to control for acoustic-related brain activity. Second, it is unknown if the individual benefit of lip movements is of behavioral relevance, as, for example, when the lips are occluded with a face mask, as has been common during the COVID-19 pandemic. Given the negative impact of face masks on behavioral measures (e.g. Rahne et al., 2021; Toscano & Toscano, 2021; Truong et al., 2021), a relationship is plausible: Individuals who benefit more should, in principle, also show poorer behavioral outcomes when no lip movements are available, as they are deprived of critical visual information.

A suitable method to obtain the individual benefit of lip movements is neural tracking (Obleser & Kayser, 2019). Besides frequency-based coherence and mutual information, temporal response functions (TRFs) have gained widespread popularity (Brodbeck & Simon, 2020; Crosse et al., 2021). TRFs typically aim to predict the M/EEG-recorded neural response to one or more stimulus features, and the prediction is correlated with the original signal to quantify neural tracking. This approach has so far extended our understanding of speech processing from acoustic features (Lalor et al., 2009) to higher-level linguistic features (Brodbeck, Hong, et al., 2018; Broderick et al., 2018; Gillis et al., 2021). Crucially, neural tracking can be used to disentangle the aforementioned intercorrelation of audiovisual speech by controlling for acoustic speech features (Gillis et al., 2022). This could reveal the “pure“ individual benefit of lip movements to neural speech tracking in audiovisual settings, which has not yet been shown.

Neural speech tracking has been proposed as an objective measure for speech intelligibility (Schmitt et al., 2022; Vanthornhout et al., 2018) along with a whole range of auditory and linguistic processes (Gillis et al., 2022). Previously, acoustic neural speech tracking has been related to behavioral measures such as speech intelligibility (Chen et al., 2023; Ding & Simon, 2013). Studies that involve visual speech features have established a relationship between the neural tracking of visual speech cues, so-called visemes, and lip-reading performance (Nidiffer et al., 2021) or lip movements and speech comprehension (Park et al., 2016). In sum, these findings strongly suggest a meaningful relationship between neural speech tracking and behavioral measures. Regarding the aspect of interindividual differences, Schubert et al. (2023) showed that the MEG-derived tendency of individuals to predict upcoming tones facilitates neural speech tracking, and this relationship generalizes to various audio-only listening situations. Here, we aim to combine both aspects by evaluating the relationship between interindividual differences and behavioral measures. In particular, we probe the behavioral relevance of the individual benefit of lip movements, especially when critical visual information is not available. Addressing this could further strengthen the case for the behavioral relevance of neural speech tracking as an objective measure of speech processing.

Here, we used MEG and an audiovisual speech paradigm with one or two speakers to investigate the benefit of lip movements and its behavioral relevance. Utilizing a state-of-the-art neural tracking framework with source-localized TRFs (see Figure 1), we show that lip movements are processed more strongly when speech is challenging. Additionally, we show that the neural tracking of lip movements is enhanced in multi speaker settings. When controlled for acoustic speech features, the obtained benefit of lip movements is, in general, more enhanced in the multi speaker condition, with substantial interindividual variability. Using Bayesian modeling, we show that acoustic speech tracking is related to behavioral measures. Crucially, we demonstrate that individuals who benefit more from lip movements show a stronger drop in performance and report a higher subjective difficulty when the mouth is occluded by a surgical face mask. In terms of neural tracking, our results suggest that individuals benefit from lip movements in a highly variable manner. We also establish a novel link between the neural benefit of visual speech and behavior when no visual speech information is available.

**Figure 1.**
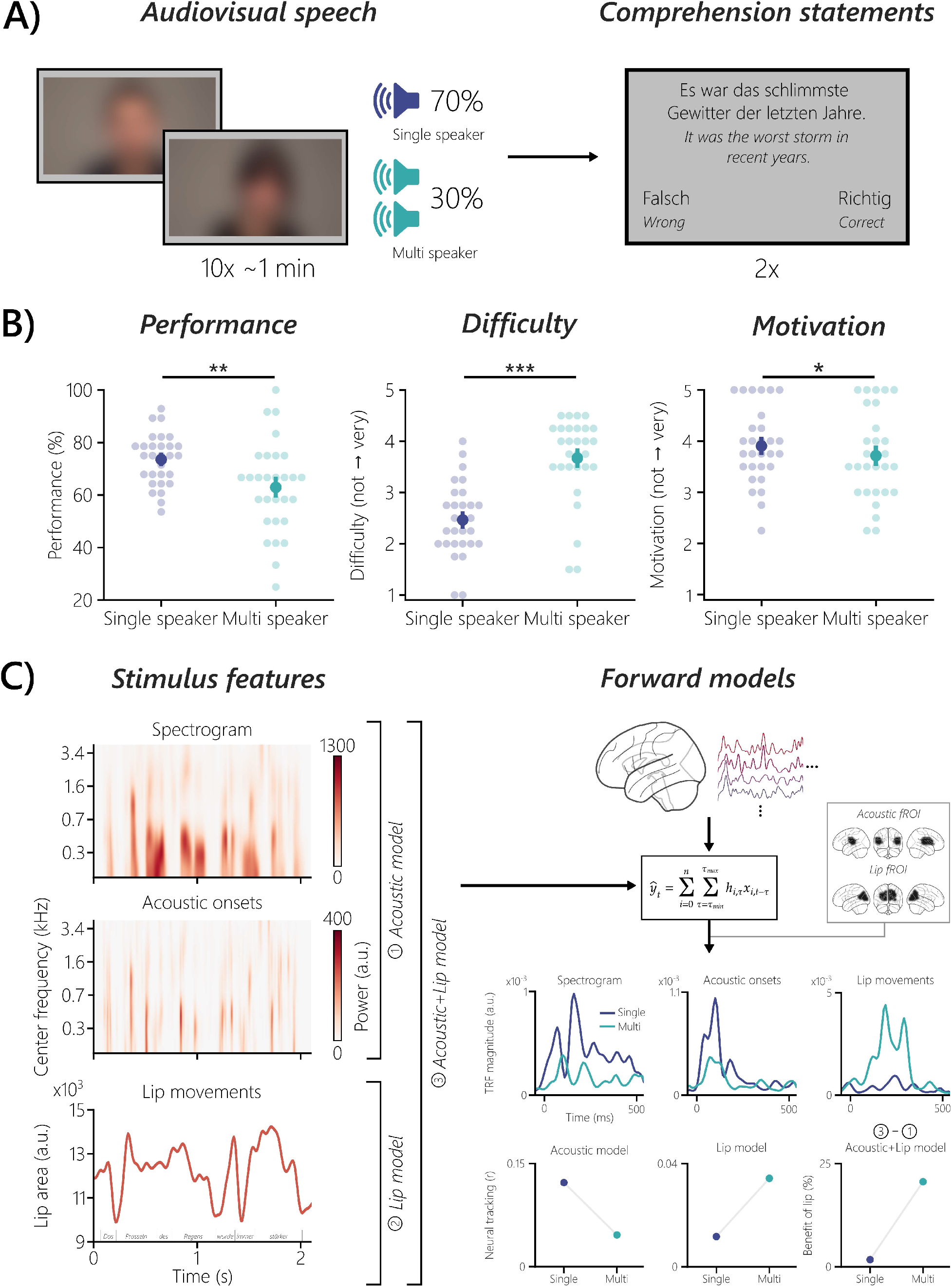
Experimental design, behavioral results and analysis framework. (A) Each block consisted of 10 ∼1-min trials of continuous audiovisual speech by either a female or male speaker (single speaker condition). In 30% of these 10 trials, a same-sex audio-only distractor speaker was added (multi speaker condition). After every block, two comprehension statements had to be rated as correct or wrong. (B) Performance on the comprehension statements in the multi speaker condition was lower than in the single speaker condition (*p* = .003, *r_C_* = 0.64). Subjective difficulty ratings, reported on a five-point Likert scale, were higher in the multi speaker condition (*p* = 9.00e^-06^, *r_C_* = -0.95). The reported motivation was lower in the multi speaker condition (p = .024, *r_C_* = 0.62). The middle dots represent the mean, and the bars, the standard error of the mean. (C) Three stimulus features (spectrogram, acoustic onsets and lip movements) extracted from the audiovisual stimuli are shown for an example sentence. Higher values in the lip area unit represent a wider opening of the mouth and vice versa. Three forward models were calculated: (1) one using only acoustic features, (2) one using only lip movements, and (3) one combining all features. Together with the corresponding source-localized MEG data, the boosting algorithm was used to calculate the models. Exemplary minimum-norm source estimates are shown for a representative participant. The resulting TRFs (a.u.) and neural tracking (expressed as Pearson’s *r*) were analyzed in functional regions of interest (fROIs), obtained either via the acoustic or lip model of the multi speaker condition. The TRFs and prediction accuracies shown are from a representative participant reflecting the group-level results. To obtain the benefit of lip movements, acoustic features were controlled by subtracting the prediction accuracies in an acoustic+lip fROI of the acoustic model from the combined model. The benefit of lip movements was expressed as a percentage change. \**p* < .05, \*\**p* < .01, \*\*\**p* < .001; *Speakers have been blurred due to a bioRxiv policy on the inclusion of faces*.

## Material and Methods

### Participants

The data was collected as part of a recent study (Haider et al., 2022), in which 30 native speakers of German participated. One participant was excluded because signal source separation could not be applied to the MEG dataset. This led to a final sample size of 29 participants (12 females, M_age_ = 26.79, SD_age_ = 4.87 years). All participants reported normal vision and hearing (thresholds did not exceed 25 dB HL at any frequency from 125 to 8000 Hz), the latter verified by a standard clinical audiometer (AS608 Basic; Interacoustics A/S, Middelfart, Denmark). Additional exclusion criteria included non-removable magnetic objects and any psychiatric or neurologic history. All participants signed an informed consent and were reimbursed at a rate of 10 € per hour. The experimental protocol was approved by the ethics committee of the Paris-Lodron-University of Salzburg and was conducted in accordance with the Declaration of Helsinki.

### Stimuli and experimental design

The experimental procedure was implemented in MATLAB 9.10 (The MathWorks Inc., Natick, Massachusetts, USA) using custom scripts. Presentation of stimuli and response collection was achieved with the Objective Psychophysics Toolbox (o_ptb; Hartmann & Weisz, 2020), which adds a class-based abstraction layer onto the Psychophysics Toolbox version 3.0.16 (Brainard, 1997; Kleiner et al., 2007; Pelli, 1997). Stimuli and triggers were generated and emitted via the VPixx system (DATAPixx2 display driver, PROPixx DLP LED projector, RESPONSEPixx response box by VPixx Technologies Inc., Saint-Bruno, Canada). Videos were back-projected onto a translucent screen with a screen diagonal of 74 cm (∼110 cm in front of the participants), with a refresh rate of 120 Hz and a resolution of 1920×1080 pixels. Timings were measured with the Black Box ToolKit v2 (The Black Box ToolKit Ltd., Sheffield, UK) to ensure accurate stimulus presentation and triggering.

The audiovisual stimuli were excerpts from four German stories, two of each read out loud by a female or male speaker (female: “Die Schokoladenvilla - Zeit des Schicksals. Die Vorgeschichte zu Band 3” by Maria Nikolai, “Die Federn des Windes” by Manuel Timm; male: “Das Gestüt am See. Charlottes großer Traum” by Paula Mattis and “Gegen den Willen der Väter” by Klaus Tiberius Schmidt). A Sony NEX-FS100 (Sony Group Corporation, Tokyo, Japan) camera with a sampling rate of 25 Hz and a RØDE NTG2 microphone (RØDE Microphones Pty. Ltd., Sydney, Australia) with a sampling rate of 48 kHz were used to record the stimuli. Each of the four stories was recorded twice, once with and once without a surgical face mask (type IIR three-layer disposable medical mask). These eight videos were cut into 10 segments of about one minute each (M = 64.29 s, SD = 4.87 s), resulting in 80 videos. In order to rule out sex-specific effects, 40 videos (20 with a female speaker and 20 with a male speaker) were presented to each participant. The speakers’ syllable rates were analyzed using Praat (Boersma, 2001; de Jong & Wempe, 2009) and varied between 3.7 Hz and 4.6 Hz (M = 4.1 Hz). The audio-only distractor speech consisted of pre-recorded audiobooks (see Schubert et al., 2023), read by either a female or a male speaker.

Before the experiment, a standard clinical audiometry was performed (for details, see *Participants*). The MEG measurement started with a 5-minute resting-state recording (not analyzed in this manuscript). Next, the participant’s individual hearing threshold was determined in order to adjust the stimulation volume. If the participant reported that the stimulation was not loud enough or comfortable, the volume was manually adjusted to the participant’s requirements.

The actual experiment consisted of four stimulation blocks, one for each of the four stories, with two featuring each sex. Each story was presented as a block of 10 ∼one-minute trials in chronological order to preserve the story content (Figure 1A). In every block, a same-sex audio-only distractor speaker was added to three randomly selected trials, with a 5-second delay and volume equal to the target speaker. The resulting ratio of 30% multi speaker trials and 70% single speaker trials per block was chosen because of a different data analysis method in Haider et al. (2022). The distractor speech started with a delay of 5 seconds to give participants time to attend the target speaker. In two randomly selected blocks, the target speaker wore a face mask (only the corresponding behavioral data was used here, see *Statistical analysis and Bayesian modeling*). Two unstandardized correct or wrong statements about semantic content were presented after each trial to assess comprehension performance and to maintain attention (Figure 1A). On four occasions in each block, participants also rated subjective difficulty and motivation on a five-point Likert scale (not depicted in Figure 1A). The participants responded by pressing buttons. The total duration of the experiment was ∼2 h, including preparation.

### MEG data acquisition and preprocessing

Before entering the magnetically shielded room, five head position indicator (HPI) coils were applied on the scalp. Electrodes for electrooculography (EOG; vertical and horizontal eye movements) and electrocardiography (ECG) were also applied (recorded data not used here). Fiducial landmarks (nasion and left/right pre-auricular points), the HPI locations and ∼300 head shape points were sampled with a Polhemus FASTRAK digitizer (Polhemus, Colchester, Vermont, USA).

Magnetic brain activity was recorded with a Neuromag Triux whole-head MEG system (MEGIN Oy, Espoo, Finland) using a sampling rate of 1000 Hz (hardware filters: 0.1-330 Hz). The signals were acquired from 102 magnetometers and 204 orthogonally placed planar gradiometers at 102 different positions. The system is placed in a standard passive magnetically shielded room (AK3b; Vacuumschmelze GmbH & Co. KG, Hanau, Germany).

A signal space separation (SSS; Taulu & Kajola, 2005; Taulu & Simola, 2006) algorithm implemented in MaxFilter version 2.2.15 provided by the MEG manufacturer was used. The algorithm removes external noise from the MEG signal (mainly 16.6 Hz, and 50 Hz, plus harmonics) and realigns the data to a common standard head position (to [0 0 40] mm, -*trans default* MaxFilter parameter) across different blocks, based on the measured head position at the beginning of each block.

Preprocessing of the raw data was done in MATLAB 9.8 using the FieldTrip toolbox (revision f7adf3ab0; Oostenveld et al., 2011). A low-pass filter of 10 Hz (hamming-windowed sinc FIR filter, onepass-zerophase, order: 1320, transition width: 2.5 Hz) was applied, and the data was downsampled to 100 Hz. Afterwards, a high-pass filter of 1 Hz (hamming-windowed sinc FIR filter, onepass-zerophase, order: 166, transition width: 2.0 Hz) was applied.

Independent component analysis (ICA) was used to remove eye and cardiac artifacts (data was filtered between 1-100 Hz, sampling rate: 1000 Hz) via the infomax algorithm (“runica” implementation in EEGLAB; Bell & Sejnowski, 1995; Delorme & Makeig, 2004) applied to a random block of the main experiment. Prior to the ICA computation, we performed a principal component analysis (PCA) with 50 components in order to ease the convergence of the ICA algorithm. After visual identification of artifact-related components, an average of 2.38 components per participant were removed (SD = 0.68).

The cleaned data was epoched into trials that matched the length of the audiovisual stimuli. To account for an auditory stimulus delay introduced by the tubes of the sound system, the data were shifted by 16.5 ms. In the multi speaker condition, the first 5 seconds of data were removed to match the onset of the distractor speech. The last eight trials were removed to equalize the data length between the single speaker and multi speaker conditions. To prepare the data for the following steps, the trials in each condition were concatenated. This resulted in a data length of ∼6 min per condition.

### Source localization

Source projection of the data was done with MNE-Python 1.1.0 running on Python 3.9.7 (Gramfort et al., 2013, 2014). A semi-automatic coregistration pipeline was used to coregister the FreeSurfer “fsaverage” template brain (Fischl, 2012) to each participant’s head shape. After an initial fit using the three fiducial landmarks, the coregistration was refined with the Iterative Closest Point (ICP) algorithm (Besl & McKay, 1992). Head shape points that were more than 5 mm away from the scalp were automatically omitted. The subsequent final fit was visually inspected to confirm its accuracy. This semi-automatic approach performs comparably to manual coregistration pipelines (Houck & Claus, 2020).

A single-layer boundary element model (BEM; Akalin-Acar & Gençer, 2004) was computed to create a BEM solution for the “fsaverage” template brain. Next, a volumetric source space with a grid of 7 mm was defined, containing a total of 5222 sources (Kulasingham et al., 2020). In order to remove non-relevant regions and shorten computation times, subcortical structures along the midline were removed, reducing the source space to 3053 sources (similar to Das et al., 2020). Subsequently, the forward operator (i.e. lead field matrix) was computed using the individual coregistrations, the BEM and the volume source space.

Afterwards, the data were projected to the defined sources using the Minimum Norm Estimate method (MNE; Hämäläinen & Ilmoniemi, 1994). MNE is known to be biased towards superficial sources, which can be reduced by applying depth weighting with a coefficient between 0.6 and 0.8 (Lin et al., 2006). For creating the MNE inverse operator, depth weighting with a coefficient of 0.8 was used (e.g. Brodbeck et al., 2018). The required noise covariance matrix was estimated with an empty-room MEG recording relative to the participant’s measurement date with the same preprocessing settings as the MEG data of the actual experiment (see *MEG data acquisition and preprocessing*). The MNE inverse operator was then applied to the concatenated MEG data with ℓ2 regularization (signal-to-noise ratio (SNR) = 3 dB, 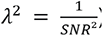) and three free-orientation dipoles orthogonally at each source.

### Extraction of stimulus features

Since the focus of this study is on audiovisual speech, we extracted acoustic (spectrograms and acoustic onsets) and visual (lip movements) speech features from the stimuli (for examples see Figure 1C). The spectrograms of the auditory stimuli were obtained using the Gammatone Filterbank Toolkit 1.0 (Heeris, 2013), with frequency cutoffs at 20 and 5000 Hz, 256 filter channels and a window time of 0.01 s. This toolkit computes a spectrogram representation on the basis of a set of Gammatone filters which are inspired by the human auditory system (Slaney, 1998). The resulting filter outputs with logarithmic center frequencies were averaged into eight frequency bands (frequencies <100 Hz were omitted; Gillis et al., 2021). Each frequency band was scaled with exponent 0.6 (Biesmans et al., 2017) and downsampled to 100 Hz, which is the same sampling frequency as the preprocessed MEG data.

Acoustic onset representations were calculated for each frequency band of the spectrograms using an auditory edge detection model (Fishbach et al., 2001). The resulting spectrograms of the acoustic onsets are valuable predictors of MEG responses to speech stimuli (Brodbeck et al., 2020; Daube et al., 2019). A delay layer with 10 delays from 3 to 5 ms, a saturation scaling factor of 30 and a receptive field based on the derivative of a Gaussian window (SD = 2 ms) were used (Gillis et al., 2021). Each frequency band was downsampled to 100 Hz.

The lip movements of every speaker were extracted from the videos with a MATLAB script adapted from Suess et al. (2022; originally by Park et al., 2016). Within the lip contour, the area, and the horizontal and vertical axis were calculated. Only the area was used for the analysis, which leads to results comparable to using the vertical axis (Park et al., 2016). The lip area signal was upsampled from 25 Hz to 100 Hz using FFT-based interpolation.

### Forward models

A linear forward modeling approach was used to predict the MEG response to the aforementioned stimulus features (see Figure 1C). These approaches are based on the idea that the brain’s response to a stimulus is a continuous function in time (Lalor et al., 2006). The boosting algorithm (David et al., 2007), implemented in eelbrain 0.38 (running on Python 3.9.7; Brodbeck et al., 2022), was used to predict MNE source-localized MEG responses to stimulus features (“MNE-boosting”; Brodbeck, Presacco, et al., 2018). For multiple stimulus features, the linear forward model can be formulated as:

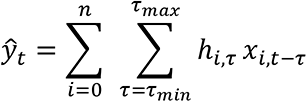

For every *n* stimulus feature, the algorithm finds an optimal filter kernel *h*, which is also known as a temporal response function (TRF). When *n* stimulus features is > 1, *h* is referred to as multivariate TRF (mTRF). The term τ denotes the delays between the predicted brain response *ŷ_t_* and stimulus feature *x* (for further details see Brodbeck et al., 2022). TRFs reflect responses to continuous data instead of averaged responses to discrete events (Crosse et al., 2021). For the estimation of the TRFs, the stimulus features and MEG data were normalized (z-scored), and an integration window from -100 to 600 ms with a kernel basis of 50 ms Hamming windows was defined. To prevent overfitting, early stopping based on the ℓ2 norm was used. By using four-fold nested crossvalidation (two training folds, one validation fold, and one test fold), each partition served as a test set once (Brodbeck et al., 2022). TRFs were estimated for each of the three free-orientation dipoles independently at all 3053 sources (see *Source localization*). The spectrogram and acoustic onset mTRFs were averaged over the frequency dimension. To account for interindividual anatomical differences, TRFs were spatially smoothed with a Gaussian kernel (SD = 5 mm; Kulasingham et al., 2020). The vector norm of the smoothed TRFs was taken, resulting in one TRF per source.

To obtain a measure of neural tracking, the predicted brain response *ŷ_t_* is correlated with the original response to calculate the prediction accuracy and computed as the average dot product over time (expressed as Pearson correlation coefficient *r*). This correlation can be interpreted as follows: The higher the prediction accuracy, the higher the neural tracking (Gillis et al., 2022).

In order to investigate the neural processing of the audiovisual speech features, we calculated three different forward models per condition and participant (see Figure 1C for the analysis framework). The acoustic model consisted of the two acoustic stimulus features (spectrogram and acoustic onsets) and – also applicable to all other models – the corresponding MNE source-localized MEG data. The lip model contained only the lip movements as a stimulus feature. Additionally, a combined acoustic+lip model was calculated to control for acoustic features in a subsequent analysis.

We defined functional regions of interest (fROIs; Nieto-Castanon et al., 2003) by creating labels based on the 90th percentile of the whole-brain prediction accuracies in the multi speaker condition (similar to Suess, Hauswald, Reisinger, et al., 2022). The multi speaker condition was chosen for extracting the fROIs because it potentially incorporates all included stimulus features, due to its higher demand (Golumbic et al., 2013). This was done separately for the acoustic and lip models to map their unique neural sources (see Figure 1C). According to the “aparc” FreeSurfer parcellation (Desikan et al., 2006), the acoustic fROI mainly involved sources in the temporal, lateral parietal and posterior frontal lobes. The superior parietal and lateral occipital lobes made up the majority of the lip fROI. To obtain an audiovisual fROI for the acoustic+lip model, we combined the labels of the acoustic and lip fROIs.

For every model, the TRFs in their respective fROI were averaged and, exclusively for Figure 2A, smoothed over time with a 50 ms Hamming window. Grand-average TRF magnitude peaks were detected with scipy version 1.8.0 (running on Python 3.9.7; Virtanen et al., 2020) and visualized as a difference between the multi and single speaker conditions. To suppress regression artifacts that typically occur (Crosse, Di Liberto, et al., 2016), TRFs were visualized between -50 and 550 ms. Prediction accuracies in the fROIs were Fisher z-transformed, then averaged, and then the z-values were back-transformed to Pearson correlation coefficients (Corey et al., 1998). For the lower panels of each model in Figure 2B, the prediction accuracies of the acoustic and lip models were averaged in their respective fROIs. Figures were created with the built-in plotting functions of eelbrain and seaborn version 0.12.0 (running on Python 3.9.7; Waskom, 2021).

**Figure 2.**
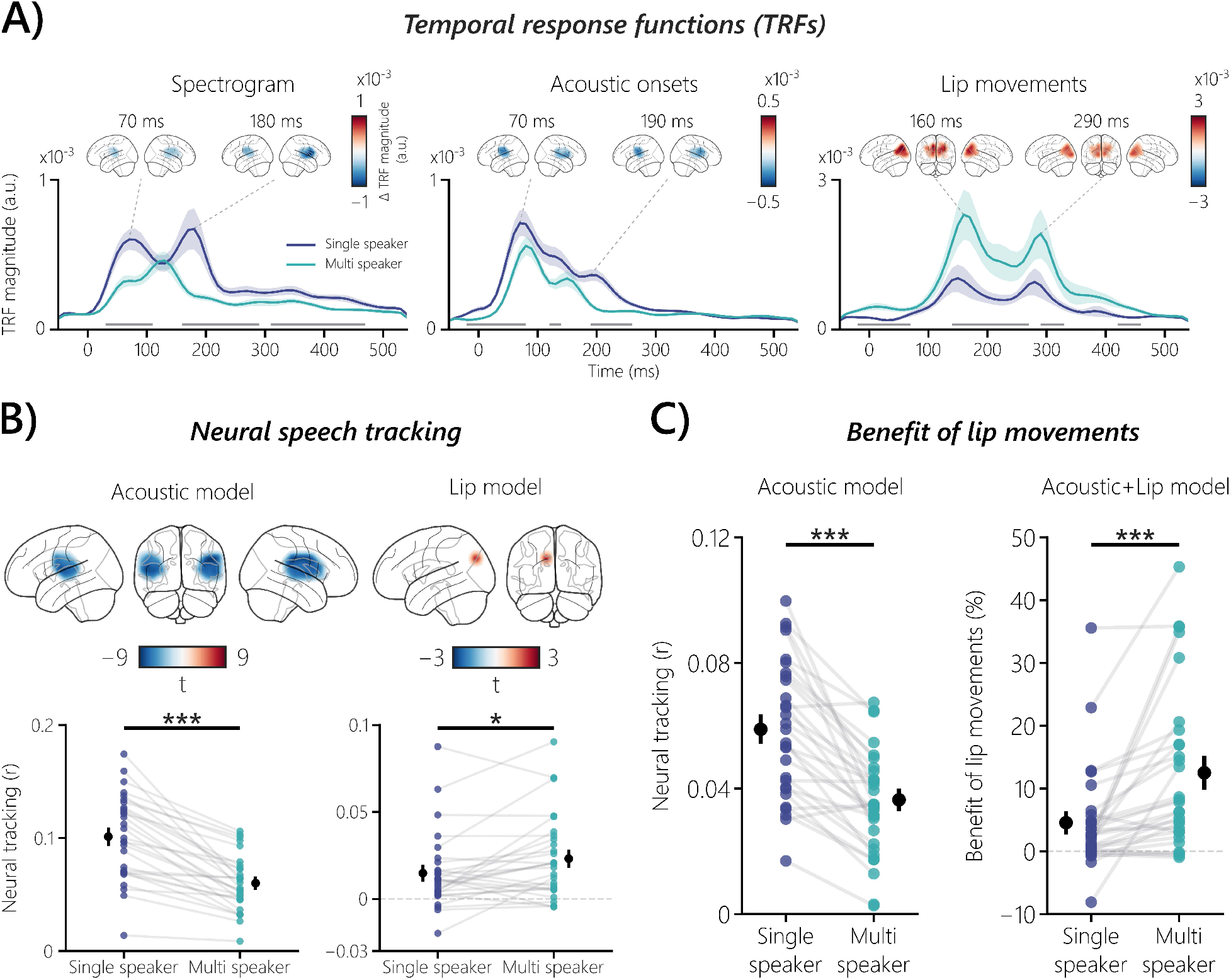
Neural responses to audiovisual speech features, neural speech tracking, and the benefit of lip movements. (A) The three plots show grand-averaged TRFs for the stimulus features in their respective fROIs and the peak magnitude contrasts (multi speaker vs. single speaker) between the two conditions in the involved sources. For the acoustic features, TRF magnitudes were generally enhanced when speech was clear, with significant differences ranging from *p* = .004 to *p* < .001 (*d* = -0.81 to -1.43). In contrast, the TRF to lip movements showed an enhanced magnitude in the multi speaker condition (*p* = .01 to *p* = .0005 and effect sizes from *d* = 0.86 to 0.91). The shaded areas of the respective conditions represent the standard error of the mean (SEM). Gray bars indicate the temporal extent of significant differences (*p* < .05) between the two conditions. (B) Neural speech tracking is shown for the non-averaged and averaged fROIs of the acoustic and lip models. Acoustic neural tracking was higher in the single speaker condition, with significant left- and right-hemispheric differences (both *p* < .001 with *d* from -1.30 to -1.47; averaged: *p* = 8.76e^-09^, *d* = -1.30). Lip movements were tracked higher in the multi speaker condition (*p* = .037, *d* = 0.51; averaged: *p* = .026, *r_C_* = 0.48). In the averaged plots, the black dots represent the mean, and the corresponding bars the SEM, of the respective condition. (C) In a combined acoustic and lip fROI, the acoustic model showed higher neural tracking in the single speaker condition (*p* = 7.68e^-08^, *d* = 1.18). The benefit of lip movements was obtained by subtracting the acoustic model from the acoustic+lip model and expressed as percentage change. Lip movements especially enhanced neural tracking in the multi speaker condition (*p* = .00003, *r_C_* = 0.89). Participants showed high interindividual variability with a visual benefit of up to 45.37%, but also only a small benefit or no benefit at all. The black dots represent the mean, and the corresponding bars the SEM, of the respective condition. \**p* < .05, \*\**p* < .01, \*\*\**p* < .001

In order to answer the question whether or not lip movements enhance neural tracking, a control for acoustic features is needed. This is particularly important due to the intercorrelation of speech features (Chandrasekaran et al., 2009; Daube et al., 2019). To investigate the individual benefit of lip movements, we used the averaged prediction accuracies in the audiovisual fROI and subtracted the acoustic model from the acoustic+lip model (for a general overview on control approaches see Gillis et al., 2022). The resulting individual benefit of lip movements was expressed as percentage change (see Figure 2C).

### Statistical analysis and Bayesian modeling

All frequentist statistical tests were conducted with built-in functions from eelbrain and the statistical package pingouin version 0.5.2 (running on Python 3.9.7; Vallat, 2018). The three behavioral measures (performance, difficulty, and motivation; Figure 1B) were statistically compared between the two conditions (single speaker and multi speaker) using a Wilcoxon signed-rank test and the matched-pairs rank-biserial correlation *r_C_*was reported as effect size (King et al., 2018).

The TRFs corresponding to the three stimulus features (spectrogram, acoustic onsets and lip movements; Figure 2A), were tested for statistical difference between the two conditions using a cluster-based permutation test with threshold-free cluster enhancement (TFCE; dependent samples t-test, 10000 randomizations, Maris & Oostenveld, 2007; Smith & Nichols, 2009). Due to the previously mentioned TRF regression artifacts, the time window for the test was limited to -50 to 550 ms. Depending on the direction of the cluster, the maximum or minimum *t*-value was reported and Cohen’s *d* of the averaged temporal extent of the cluster was calculated.

We tested the non-averaged prediction accuracies in the acoustic and lip fROIs (Figure 2B) with a cluster-based permutation test with TFCE (dependent samples t-test, 10000 randomizations). According to the cluster’s direction, the maximum or minimum *t*-value was reported, and Cohen’s *d* of the cluster’s averaged spatial extent was calculated. Additionally, averaged prediction accuracies in the acoustic and lip fROIs were statistically tested with a dependent-samples t-test, and Cohen’s *d* was reported as effect size. In the audiovisual fROI, the prediction accuracies and benefit of lip movements (Figure 2C) were tested with a dependent-samples t-test, and Cohen’s *d* was reported as effect size. If the data were not normally distributed according to a Shapiro-Wilk test, the Wilcoxon signed-rank test was used, and the matched-pairs rank-biserial correlation *r_C_* was reported as effect size. The distribution of the benefit of lip movements was assessed using the bimodality coefficient (Freeman & Dale, 2013).

To investigate if neural tracking is predictive for behavior, we calculated Bayesian multilevel models in R version 4.2.2 (R Core Team, 2022) with the Stan-based package brms version 2.18.4 (Bürkner, 2017; Carpenter et al., 2017). Neural tracking (i.e. the averaged prediction accuracies within the respective fROI) was used to separately predict the three behavioral measures. A random intercept was added for each participant to account for repeated measures (single speaker and multi speaker). The models were fitted independently for the acoustic and lip models (Figure 3). According to the Wilkinson notation (Wilkinson & Rogers, 1973), the general formula was:

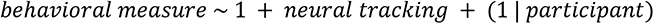

**Figure 3.**
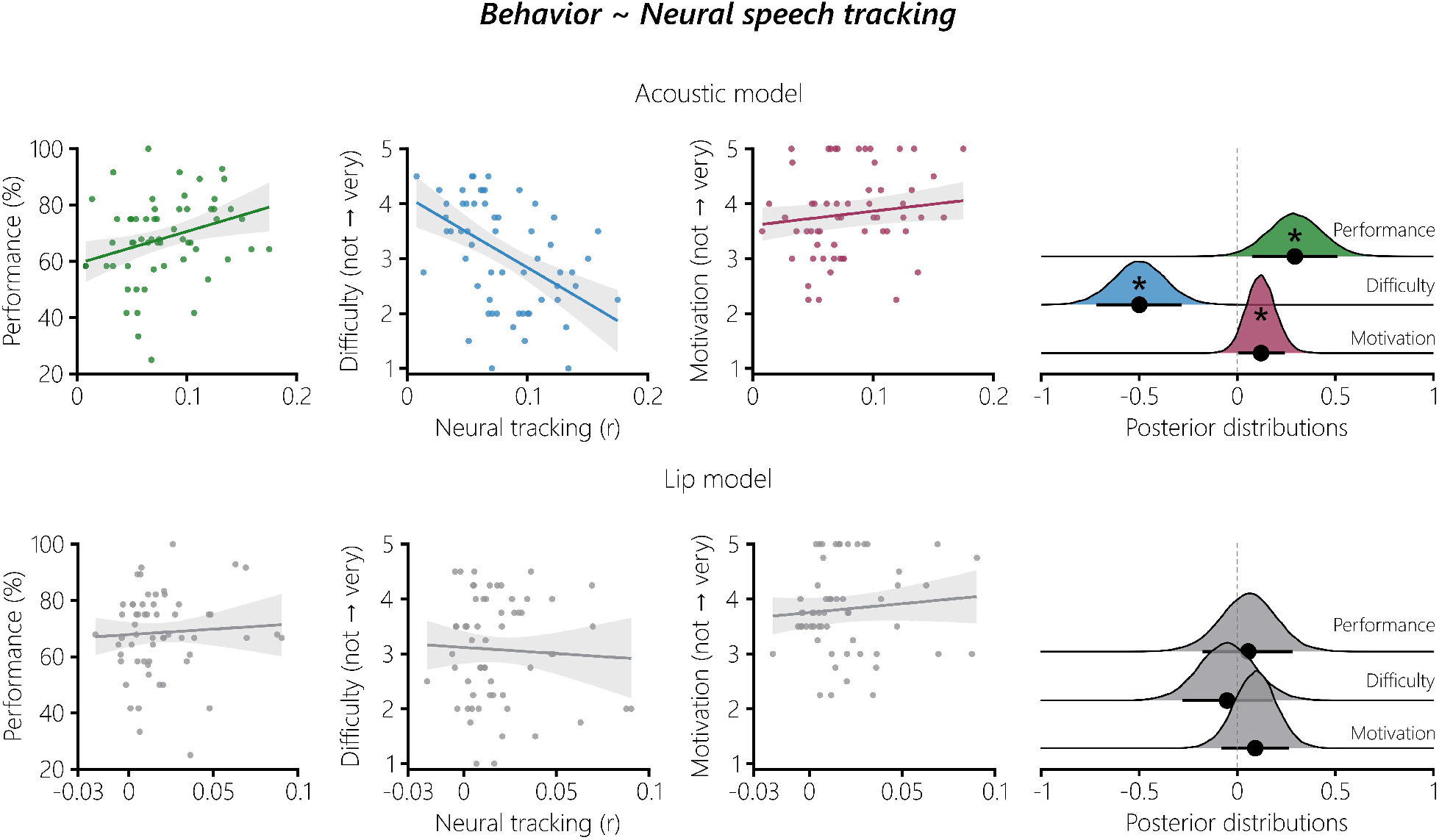
Relating behavior to neural speech tracking. Bayesian multilevel models were fitted to predict the behavioral measures with neural speech tracking. Higher acoustic neural speech tracking was linked to higher performance, lower difficulty ratings and higher motivation ratings. No evidence for an effect was observed for the neural tracking of lip movements. The shaded areas show the 89% CIs of the respective model. The distributions on the right show the posterior draws of the three models. The black dots represent the mean standardized regression coefficient *b* of the corresponding model. The corresponding bars show the 89% CI. If zero was not part of the 89% CI, the effect was considered significant (*).

We wanted to test whether the individual benefit of lip movements to neural speech tracking (see *Forward models*) yields any behavioral relevance. For this, we also used the behavioral data of the otherwise unanalyzed conditions with a face mask (see *Stimuli and experimental design*). We fitted Bayesian multilevel models with the individual benefit of lip movements to separately predict the behavioral measures when the speaker wore a face mask or not (Figure 4). The general formula was:

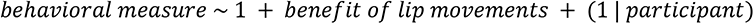

**Figure 4.**
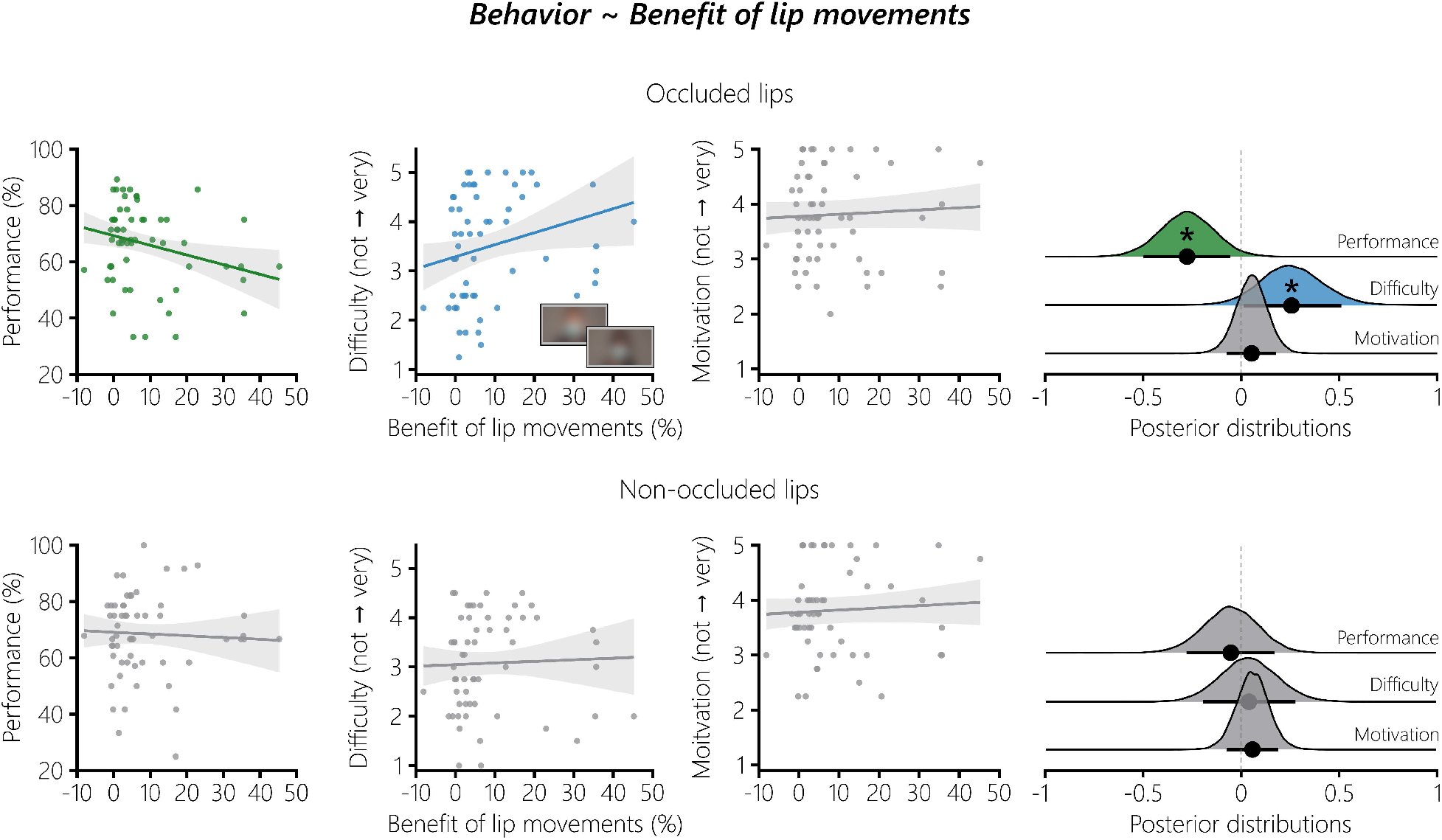
Relating the benefit of lip movements to behavior. The benefit of lip movements was used to predict the behavioral measures when the lips are occluded or not. The values of the fitted Bayesian multilevel models are shown with a depiction of the conditions in which the speakers wore a surgical face mask. When the benefit of lip movements was high, performance was lower, and difficulty was reported higher. No evidence for an effect was observed for the motivation rating. The behavioral measures when the lips were not occluded were not linked to the benefit of lip movements. The shaded areas show the 89% CIs of the respective model. The distributions on the right show the posterior draws of the three models. The black dots represent the mean standardized regression coefficient *b* of the corresponding model. The corresponding bars show the 89% CI. If zero was not part of the 89% CI, the effect was considered significant (*). *Speakers have been blurred due to a bioRxiv policy on the inclusion of faces*.

Before doing so, we fitted control models to show the effect of the conditions on the behavioral measures when the lips are occluded (see Supplementary Table 1). Additional control models to test the effect of the benefit of lip movements on the behavioral data without a face mask were also fitted (see Supplementary Table 2 for model fits). In all described models, a random intercept was included for each participant to account for repeated measures (single speaker and multi speaker).

Weakly or non-informative default priors of brms were used, whose influence on the results is negligible (Bürkner, 2017, 2018). For model calculation, all numerical variables were z-scored, and standardized regression coefficients (*b*) were reported with 89% credible intervals (CIs; i.e. Bayesian uncertainty intervals, McElreath, 2020). In addition, we report posterior probabilities (PP*_b_*_>0_) with values closer to 100%, providing evidence that the effect is greater than zero, and closer to 0% that the effect was reversed (i.e. smaller than zero). If the 89% CIs for an estimate did not include zero and PP*_b_*_>0_ was below 5.5% or above 94.5%, the effects were considered statistically significant.

All models were fitted with a Student-t distribution, as indicated by graphical posterior predictive checks, Pareto *k^* diagnostics (Vehtari, Simpson, et al., 2022) and leave-one-out crossvalidation via loo version 2.5.1 (Vehtari et al., 2017; Vehtari, Gabry, et al., 2022). Common algorithm-agnostic (Vehtari et al., 2021) and algorithm-specific diagnostics (Betancourt, 2018) showed that all Bayesian multilevel models converged. For all relevant parameters, the convergence diagnostic R^ < 1.01 and effective sample size (ESS) > 400 indicated that there were no divergent transitions. Figures were created with ggplot2 version 3.4.0 (Wickham, 2016) and ggdist version 3.2.0 (Kay, 2022). Unstandardized *b*’s were used for the fitted values of the models in Figures 3 and 4.

### Data and Code Availability

Preprocessed data and code are publicly available at GitHub (https://github.com/reispat/ av_speech_mask).

## Results

Twenty-nine participants listened to audiobooks with a corresponding video of the speaker and a randomly occurring audio-only distractor. Source-localized MEG responses to acoustic features (spectrogram and acoustic onsets) and lip movements were predicted using forward models (TRFs). We compared the TRFs between the two conditions and evaluated neural tracking of the acoustic features and lip movements. The individual benefit of lip movements was obtained by controlling for acoustic features and was compared between conditions. Using Bayesian multilevel modeling, we predicted the behavioral measures with neural tracking. We also probed the individual benefit of lip movements for their behavioral relevance by predicting the behavioral measures when the lips were occluded with a surgical face mask or not.

### Listening situations with multiple speakers are behaviorally more demanding

Participants performed worse in the multi speaker condition (M = 62.93%, SD = 17.34%), compared to the single speaker condition (M = 73.52%, SD = 9.71%; *W* = 73.00, *p* = .003, *r_C_* = 0.64). In the multi speaker condition, subjective difficulty ratings were higher (M = 3.67, SD = 0.82) than in the single speaker condition (M = 2.47, SD = 0.71; *W* = 11.50, *p* = 9.00e^-06^, *r_C_* = -0.95). Motivation was rated higher in the single speaker condition (M = 3.91, SD = 0.74) compared to the multi speaker condition (M = 3.72, SD = 0.85; *W* = 29.00, *p* = .024, *r_C_* = 0.62). Overall, behavioral data showed that in the multi speaker condition, participants performed worse, reported the task to be more difficult and were less motivated (Figure 1B).

### Neural responses to lip movements are enhanced in a multi speaker setting

First, we analyzed the neural responses to acoustic and visual speech features by statistically comparing the corresponding TRFs between the single- and multi speaker conditions within their respective fROIs (Figure 2A). The spectrogram TRFs showed a significant difference between conditions, with three clusters extending from early (30 to 110 ms; *t =* -5.26, *p* = .0001, *d* = -0.81), middle (160 to 290 ms; *t* = -3.78, *p* = .003, *d* = -1.00) and late (310 to 470 ms; *t* = -5.58, *p* = .0001, *d* = -1.02) time ranges. Grand-average TRF peaks are more pronounced in the single speaker condition, with two peaks at 70 and 180 ms. While the first peak is also present in the multi speaker condition, the second peak appeared 50 ms earlier than the single speaker setting. The latter peak caused the largest differences in the magnitudes of the TRFs, which are most prominent in the right hemisphere of the fROI.

The TRFs to acoustic onsets showed a significant difference between single- and multi speaker speech, with three clusters extending from early (-20 to 80 ms; *t* = -5.39, *p* < .001, *d* = -1.10; Figure 2A), mid (120 to 140 ms; *t* = -4.54, *p* = .004, *d* = -1.43) and mid-late (190 to 260 ms; *t* = - 6.11, *p* < .001, *d* = -1.13) time windows. The TRFs showed two peaks at 70 and 190 ms in the single speaker condition. Similar to the spectrogram TRFs, the first peak in the multi speaker condition is at the same time point as in the single speaker condition and the second peak is 50 ms earlier. The magnitude differences across peaks and hemispheres are not substantially different.

TRFs to lip movements show an opposite pattern to the TRFs to acoustic features, with stronger processing in the multi speaker condition. Significant condition differences in the TRFs to lip movements between single- and multi speaker speech were found, with four clusters extending from early (-20 to 70 ms; *t* = 4.41, *p* = .0005, *d* = 0.86; Figure 2A), mid (140 to 270 ms; *t* = 3.97, *p* = .001, *d* = 0.88), mid-late (290 to 330 ms; *t* = 3.34, *p* = .01, *d* = 0.91) and late (420 to 460 ms; *t* = 3.90, *p* = .002, *d* = 0.90) time windows. The latencies of the peaks were later in general (160 and 290 ms), as compared to the acoustic TRFs, which is also in line with the longer duration for a stimulus to reach the visual system (Thorpe et al., 1996; VanRullen & Thorpe, 2001). In the single speaker condition, they are delayed by 10 ms, and magnitude differences are most prominent in the first peak and left hemisphere.

Our initial analysis showed that neural responses to acoustic features are stronger when speech is clear. In contrast, neural responses to lip movements were enhanced in a multi speaker environment. The stronger processing of lip movements suggests a greater reliance on the lips of a speaker when speech is harder to understand.

### The cocktail party diametrically affects acoustic and visual neural speech tracking

So far, the TRF results indicate a stronger neural response to lip movements and a weaker one to acoustic features when there is more than one simultaneous speaker. We also wanted to answer the question whether neural tracking of audiovisual speech features differs between the single speaker and multi speaker conditions in their respective fROIs (Figure 2B; see Figure S1 for whole-brain neural tracking of the audiovisual speech features). Acoustic neural tracking in the non-averaged acoustic fROI showed a significant condition difference in the left (*t* = -8.04, *p* < .001, *d* = -1.47) and right (*t* = -9.26, *p* < .001, *d* = -1.30) hemispheres. Averaged acoustic neural tracking was higher in the single speaker condition than in the multi speaker condition (*t*(28) = - 8.07, *p* = 8.76e^-09^, *d* = -1.30). Neural tracking of lip movements showed a significant condition difference in the left hemisphere (*t* = 3.83, *p* = .037, *d* = 0.51; Figure 2B), with a focal inferior parietal area involved. When averaging over sources, neural tracking was higher in the multi speaker condition than in the single speaker condition (*W* = 114.00, *p* = .026, *r_C_* = 0.48).

Overall, the results showed that neural tracking was enhanced for acoustic features when speech is clear, and higher for lip movements when there are multiple speakers. This is in line with the observed neural responses.

### Lip movements enhance neural speech tracking more in multi speaker situations

When there are two speakers, we have so far demonstrated that lip movements are processed more strongly and lead to higher neural tracking compared to one speaker. However, their unique contribution to neural tracking is still unknown, due to the intercorrelation of speech features (Chandrasekaran et al., 2009; Daube et al., 2019). To address this, we controlled for the acoustic features so as to obtain the unique benefit of lip movements over and above acoustic speech features. First, the acoustic model was evaluated in the audiovisual fROI (Figure 2C). Acoustic neural tracking was higher in the single speaker condition than in the multi speaker condition (*t*(28) = -7.20, *p* = 7.68e^-08^, *d* = 1.18). The acoustic model served as a baseline and was subtracted from a combined acoustic+lip model and expressed as percentage change. The obtained benefit of lip movements was higher in the multi speaker condition than in the single speaker condition (*W* = 24.00, *p* = .00003, *r_C_* = 0.89). The benefit of lip movements showed high interindividual variability and seemed to follow a bimodal distribution (Figure 2C), which was confirmed by a bimodality coefficient of 0.68 (values > 0.555 indicate bimodality; Pfister et al., 2013).

These results strongly indicate that lip movements enhance neural tracking, especially in multi-talker speech. However, substantial interindividual variability was observed, with participants showing an individual benefit of lip movements of up to 45.37% in the multi speaker condition, while others showed only a small benefit or no benefit at all. In the next steps, we will probe the behavioral relevance of the benefit that lip movements provide to neural speech tracking by depriving individuals of this source of information.

### Only acoustic neural speech tracking predicts behavior

Having established that listening situations with two speakers affect neural tracking of acoustic and visual speech features in a diametrical way, we were further interested if neural tracking is able to predict the behavioral measures. We calculated Bayesian multilevel models to predict the three behavioral measures (performance, difficulty and motivation; see Figure 1B) with the averaged neural tracking of the acoustic and lip models (Figure 3). In the acoustic model, higher neural tracking was linked to higher performance (*b* = 0.29, 89% CI = [0.07, 0.51], PP*_b_*_>0_ = 98.37%). Lower neural tracking predicted higher difficulty ratings (*b* = -0.50, 89% CI = [-0.72, - 0.29], PP*_b_*_>0_ = 0.01%). When neural tracking was high, the motivation ratings were also higher (*b* = 0.12, 89% CI = [0.004, 0.24], PP*_b_*_>0_ = 95.05%).

Neural tracking of lip movements was not related to performance (*b* = 0.06, 89% CI = [-0.18, 0.28], PP*_b_*_>0_ = 65.61%; Figure 3). We also observed no evidence for an effect of the difficulty (*b* = -0.05, 89% CI = [-0.28, 0.18], PP*_b_*_>0_ = 35.63%) or motivation (*b* = 0.09, 89% CI = [-0.08, 0.26], PP*_b_*_>0_ = 80.40%) ratings.

These results indicate that acoustic neural speech tracking predicts behavior: The higher the neural speech tracking, the higher the performance and motivation ratings. Lower acoustic neural speech tracking was linked to higher difficulty ratings. In contrast, neural speech tracking of lip movements did not predict behavior.

### Stronger benefit of lip movements predicts behavioral deterioration when lips are occluded

Given the finding that lip movements enhance neural speech tracking (Figure 2C), we were interested in whether this visual benefit is behaviorally relevant. To do so, we also used the behavioral data from the otherwise unanalyzed conditions in which the mouth was occluded by a surgical face mask (see Figure 4 for an example). Given that critical visual information is missing in these conditions, individuals who show a strong benefit of lip movements on a neural level should show poorer behavioral outcomes. An initial analysis showed that the effect of the conditions with a surgical face mask on behavior followed the same pattern as those with non-occluded lips (see Figure 1B), although with no effect on the motivation ratings. These control models are reported in Supplementary Table 1.

While the effects on a solely behavioral level seem not to differ when the lips are occluded or not, predicting the behavioral measures with the lip benefit showed the expected outcome (Figure 4): Participants that had a higher benefit of lip movements in terms of neural tracking showed a decline in performance (*b* = -0.27, 89% CI = [-0.49, -0.06], PP*_b_*_>0_ = 2.21%) and reported the task to be more difficult (*b* = 0.25, 89% CI = [0.01, 0.51], PP*_b_*_>0_ = 95.41%). The motivation ratings did not yield an effect (*b* = 0.05, 89% CI = [-0.07, 0.18], PP*_b_*_>0_ = 76.14%).

Interestingly, we were not able to establish a link between the benefit of lip movements to the behavioral data when the lips were not occluded (Figure 4; see Supplementary Table 2 for model fits). Taken together, these findings support a behavioral relevance of the benefit of lip movements. Individuals that benefit more from lip movements on a neural level performed worse and reported the task to be more difficult when the mouth of the speaker was covered by a surgical face mask.

## Discussion

Neural speech tracking is widely used to study the neural processing of continuous speech, though primarily with audio-only stimuli (Brodbeck, Hong, et al., 2018; Chalas et al., 2022; Di Liberto et al., 2015; Keitel et al., 2018). Recent studies have used audiovisual speech settings, but without directly modeling the visual speech features (Crosse, Liberto, et al., 2016; Golumbic et al., 2013) or not incorporating their temporal dynamics due to the use of frequency-based methods (Aller et al., 2022; Bröhl et al., 2022; Park et al., 2016). Here, we show, for the first time, the temporal dynamics and cortical origins of TRFs obtained from lip movements in an audiovisual setting with one or two speakers. Using these neural responses, we demonstrate that the neural tracking of lip movements is enhanced in a multi speaker situation compared to a single speaker. When controlling for acoustic speech features, we show that the obtained benefit of lip movements is enhanced in the multi speaker condition, although with high interindividual variability. Using Bayesian modeling, we demonstrate that acoustic neural speech tracking predicts the behavioral measures. Furthermore, individuals who displayed a higher benefit of lip movements showed a stronger behavioral decline when the mouth was occluded with a surgical face mask. Our findings show that individuals vary highly in their visual speech benefit and provide new insights into the behavioral relevance of neural speech tracking.

### Neural responses to audiovisual speech

Similar to Brodbeck, Hong, et al. (2018), neural responses to acoustic features in the two-speaker paradigm were generally weaker. The TRFs to lip movements showed an opposite pattern, with an enhanced magnitude in the multi speaker condition (Figure 2A), and with substantially later peaks compared to the TRF to acoustic features. This is in line with Bourguignon et al. (2020), where initial TRF peaks at 115 and 159 ms were shown from two significant sources, overlapping with our involved parietal and occipital sources (see Figure 1C). However, the TRFs in their work were modeled to lip movements from silent videos, which precludes a comparison between different listening situations. Our findings also strengthen the argument that TRFs to visual speech are qualitatively different from TRFs to acoustic speech (for coherence, see Park et al., 2016), despite the high intercorrelation of speech features (Chandrasekaran et al., 2009).

### Neural tracking of audiovisual speech

Based on the source-localized neural tracking, we determined fROIs via a data-driven approach – separately for the acoustic features and lip movements (see Figure 1C). The fROIs for the acoustic speech features involved sources along temporal, parietal and posterior frontal regions, covering regions that are related to speech perception (Franken et al., 2022). Previous studies source-localized TRFs in audio-only settings, though commonly restricting the analysis to temporal regions (e.g. Brodbeck, Hong, et al., 2018; Kulasingham et al., 2020). The fROIs for the lip movements involved parietal and occipital regions, in line with previous studies that source-localized the neural tracking of lip movements (Aller et al., 2022; Bourguignon et al., 2020; Hauswald et al., 2018). Similar to Park et al. (2016), we also observed neural tracking of lip movements in temporal regions (see Figure S1), but with less involvement of the primary visual cortex and prominent only in the single speaker condition. Due to our approach of defining our fROIs based on the multi speaker condition, we removed any involvement of auditory regions in the lip fROIs. In contrast to Park et al. (2016), we did not observe neural tracking of lip movements in motor regions, resulting in no involvement of related sources in the lip fROIs.

When analyzing neural speech tracking in the acoustic fROIs, we showed a large effect with enhanced tracking in the single speaker condition compared to the multi speaker condition (Figure 2B). We did not find a previous study that showed such a statistical contrast, which could be due to the general focus on neural tracking of attended versus unattended speech, especially to decode auditory attention (e.g. Ciccarelli et al., 2019; Geirnaert et al., 2021; Mirkovic et al., 2015; J. A. O’Sullivan et al., 2015; Schäfer et al., 2018). On a group level, the neural tracking of lip movements showed an enhancement in the multi speaker condition (Figure 2B). When comparing the involved sources of the corresponding lip fROI, we found a medium effect in the left superior parietal cortex. This is well in line with Park et al. (2016), showing an effect in left occipital and parietal cortex when comparing two similar conditions to our design (“AV congruent vs. All congruent”), although after partializing out auditory-related coherence. When we averaged the neural tracking of lip movements, we observed interesting patterns, with participants showing no meaningful neural tracking (i.e. close to zero or negative correlations) when there was one speaker, but when speech became challenging, their neural tracking reached positive values. Notably, this pattern was reversed for some participants, suggesting that not all of them used the lip movements in the same manner. To investigate this further, eye tracking should be used to identify which face regions participants fixated when attending audiovisual speech (e.g. Rennig & Beauchamp, 2018) or to additionally incorporate a recently proposed phenomenon termed “ocular speech tracking” (Gehmacher et al., 2023). Altogether, this is the first time that neural tracking of lip movements has been quantified in the context of TRFs, although with substantially smaller correlations as compared to acoustic speech tracking. Other algorithms, such as ridge regression, could, in principle, yield higher values due to their optimization towards maximizing neural tracking values (for a comparison of algorithms, see Kulasingham & Simon, 2022).

### Benefit of lip movements

We first compared the neural tracking of audiovisual speech between single speaker and multi speaker conditions in an isolated manner. Due to the aforementioned inter-correlation of speech features (Chandrasekaran et al., 2009; Daube et al., 2019), this approach could not rule out any acoustic contributions to the neural tracking of lip movements or vice versa. To reveal the unique benefit of lip movements and to incorporate regions that are part of models of audiovisual speech perception (Bernstein & Liebenthal, 2014) and multisensory integration (Peelle & Sommers, 2015), we combined both fROIs and controlled for acoustic speech features. Within the TRF framework, we provide first evidence that lip movements enhance acoustic-controlled neural speech tracking (Figure 2C). A general enhancement was observed for both single- and multi speaker speech, which is in line with behavioral findings that visual speech features enhance intelligibility under clear speech conditions as well (Blackburn et al., 2019; Stacey et al., 2016). When comparing the two conditions, we observed a large effect, showing a higher benefit of lip movements in the multi speaker condition. Our findings are also well in line with a previous study (Park et al., 2016) that used partial coherence to remove auditory-related contributions, showing higher coherence in a challenging audiovisual speech situation compared to a condition where the audiovisual input was congruent.

Analogous to behavioral findings in Aller et al. (2022), the benefit of lip movements showed high interindividual variability (see Figure 2C) and followed a bimodal distribution. Some individuals benefited massively from lip movements, while others showed only a small benefit or none at all. Interestingly, one individual even showed a negative influence when adding lip movements to the acoustic model when there was only one speaker. As soon as speech became challenging, that individual benefited from the lip information. Overall, these findings are in line with the beneficial effects of visual speech when listening is challenging (e.g. Grant & Seitz, 2000; Remez, 2012; Ross et al., 2007; Sumby & Pollack, 1954). Given our moderate sample size, we refrained from conducting further analysis by defining groups of individuals who showed a higher or lower benefit of lip movements. Future studies should include more participants, as well as hearing-impaired populations. A recent study that used neural tracking showed an increased audiovisual speech benefit when speech was noisy (Puschmann et al., 2019). This could also provide a clearer picture of how individuals benefit from lip movements in terms of neural tracking. Previous studies used only the acoustic envelope to investigate the benefit of visual speech features on neural speech tracking (Crosse, Liberto, et al., 2016; Golumbic et al., 2013). Here, we also incorporated lip movements to provide a more complete picture of the unique benefit of visual speech features in audiovisual settings with naturalistic stimuli (Hamilton & Huth, 2020; A. E. O’Sullivan et al., 2019).

### Predicting behavior with neural tracking

Our initial analysis of the behavioral measures suggests a higher cognitive demand when speech was challenging (Figure 1B). Participants displayed lower task performance, higher difficulty ratings and lower motivation ratings when more than one speaker was involved (Figure 1B). The influence of challenging speech is also reflected in the findings of neural speech tracking (Figure 2B). Building on these results, we used Bayesian multilevel modeling to establish a link between neural speech tracking and behavior (Figure 3). Higher acoustic neural tracking is related to higher task performance, a finding also reported in a study that used vocoded speech (Chen et al., 2023). We also show that higher acoustic neural tracking is related to lower difficulty ratings. This is in line with a study that showed a positive relationship between speech intelligibility ratings and acoustic neural tracking, though using speech-in-noise (Ding & Simon, 2013). Higher motivation ratings were associated with higher acoustic neural tracking – in contrast to Schubert et al. (2023) – showing no relationship between the two measures. We were not able to establish any link between the neural tracking of lip movements and the behavioral measures. It is important to note here that the analyzed neural tracking of lip movements was not yet controlled for speech acoustics (Gillis et al., 2022), which could confound any relationship with behavior. A recent MEG study impressively showed that the neural tracking of acoustic speech features can explain cortical responses to higher-order linguistic features, such as phoneme onsets (Daube et al., 2019), emphasizing the importance of controlling acoustics (see also Gillis et al., 2021).

The COVID-19 pandemic established the use of face masks on a global scale (Feng et al., 2020). However, it has been demonstrated that covering the mouth has adverse effects on behavioral measures, such as speech perception (e.g. Rahne et al., 2021). On a neural level, Haider et al. (2022) showed that surgical face masks impair the neural tracking of acoustic and higher-order segmentational speech features. However, the consequences of an absence of visual speech were not analyzed in this study. Here, we establish a relationship between behavioral measures and the individual benefit of visual speech on neural tracking. When the speaker wore a surgical face mask, individuals that benefit more from lip movements displayed lower task performance and higher difficulty ratings. Strikingly, no effect was found when the speaker did not wear a surgical face mask. Overall, our results suggest that individuals who use lip movements more effectively show behavioral deterioration when visual speech is absent. However, further studies with larger sample sizes are needed to disentangle the potential influence of experimental conditions on this relationship, e.g. using Bayesian mediation analysis (Nuijten et al., 2015; Yuan & MacKinnon, 2009).

### Conclusion

The current study provides first evidence for the substantial interindividual variability in the neural tracking of lip movements and its relationship to behavior. First, we show that neural responses to lip movements are more pronounced when speech is challenging, compared to when speech is clear. We show that lip movements effectively enhance neural speech tracking in brain regions related to audiovisual speech, with high interindividual variability. Furthermore, we demonstrate that this individual visual benefit is behaviorally relevant. Individuals that benefit more from lip movements have a lower task performance and rate the task to be more difficult when the speaker wears a surgical face mask. Remarkably, this relationship is completely absent when the speaker did not wear a mask. Our results provide new insights into the individual differences in the neural tracking of lip movements and offer potential implications for future clinical and audiological settings to objectively assess audiovisual speech perception.

## Acknowledgments

P.R. is supported by the Austrian Science Fund (FWF; Doctoral College “Imaging the Mind”; W 1233-B), as well as N.S. (“Audiovisual speech entrainment in deafness”; P31230) and C.H. (“Impact of face masks on speech comprehension”; P34237). P.R. is also supported by the Austrian Research Promotion Agency (FFG; BRIDGE 1 project “SmartCIs”; 871232). M.G. is supported by a Strategic Basic research grant by the Research Foundation Flanders (FWO, Grant No. 1SA0620N). J.V. is supported by a postdoctoral grant provided by the FWO (Grant No. 1290821). The authors would like to thank Juliane Schubert for stimulus recordings and Sarah Danböck for her helpful methodological input.

## Author Contributions

P.R. analyzed the data, created the figures and wrote the manuscript. M.G. and J.V. analyzed the data and edited the manuscript. N.S. and T.H. provided input on data analysis and edited the manuscript. C.H. designed the study, collected the original dataset and edited the manuscript. A.H. designed the study and edited the manuscript. K.S. edited the manuscript. T.F. supervised the project and edited the manuscript. N.W. acquired the funding, supervised the project and edited the manuscript.

## Conflict of interest statement

K.S. is an employee of MED-EL GmbH. All other authors declare no competing interests.

---

**Figure S1.**
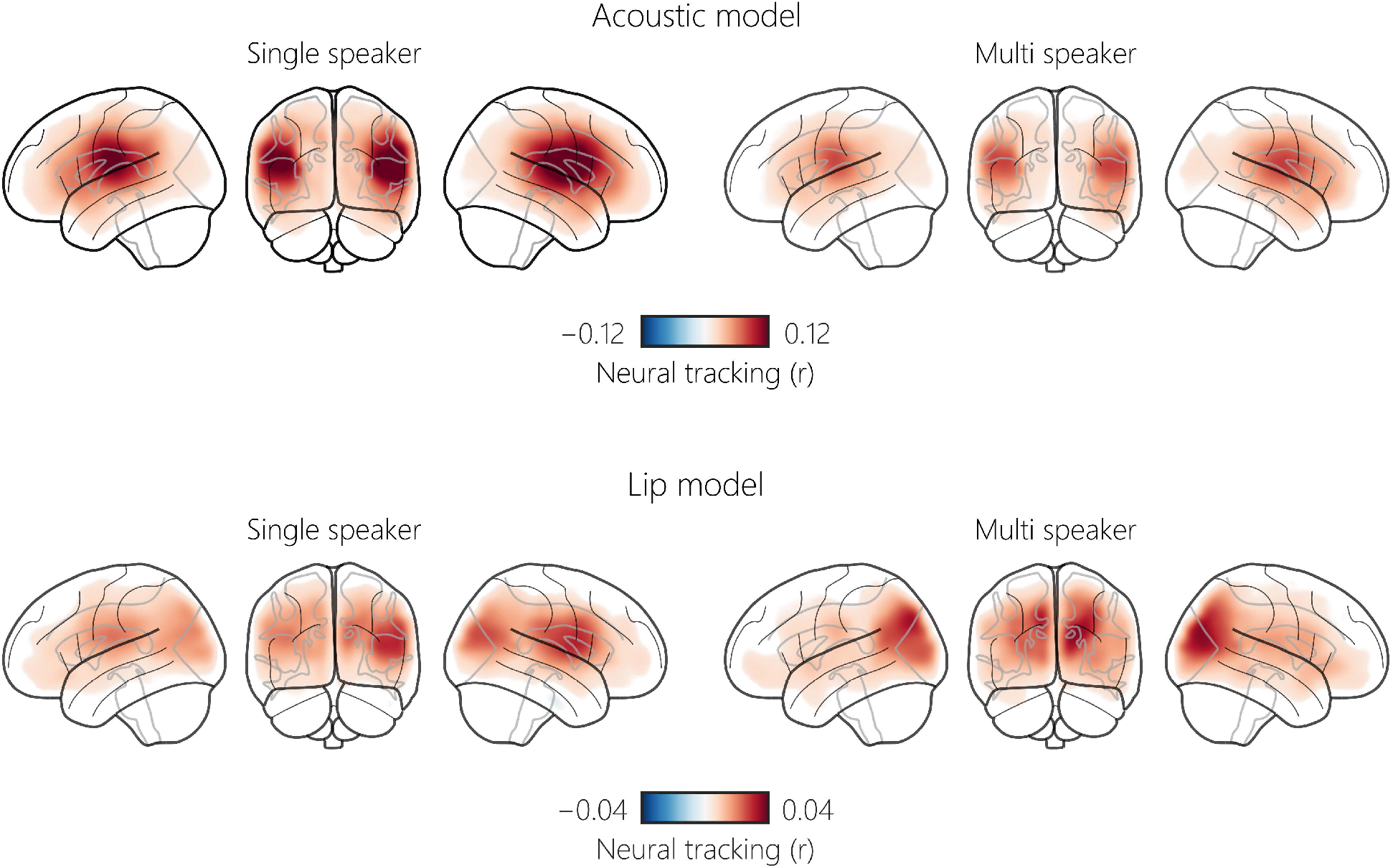
Whole-brain neural tracking of the audiovisual speech features. Neural tracking (*r*) of all sources is shown for the acoustic model (spectrogram and acoustic onsets) and the lip model (lip movements).

**Supplementary Table 1.** Effects of conditions on behavior when the lips are occluded. The formula was: behavioral measure ∼ 1 + conditions + (1 | participant). *89% CI not including zero and PP_b>0_ below 5.5% or above 94.5% (i.e. significant effect).

|  | Conditions |  |  |
| --- | --- | --- | --- |
| | $b$ | 89% CI | PP<sub>b>0</sub> |
| Performance (occluded lips) | -0.77 | [-1.13, -0.41]* | 0.07%* |
| Difficulty (occluded lips) | 1.26 | [1.04, 1.49]* | 100%* |
| Motivation (occluded lips) | -0.11 | [-0.27, 0.04] | 11.11% |

**Supplementary Table 2.** Effects of the benefit of lip movements on behavior when the lips are not occluded.

|  | Benefit of lip movements |  |  |
| --- | --- | --- | --- |
|  | *b* | 89% CI | PP<sub>b>0</sub> |
| Performance | -0.05 | [-0.28, 0.17] | 36.09% |
| Difficulty | 0.04 | [-0.19, 0.28] | 60.86% |
| Motivation | 0.06 | [-0.08, 0.19] | 76.64% |

